# Disruption of The Psychiatric Risk Gene Ankyrin 3 Enhances Microtubule Dynamics Through GSK3/CRMP2 Signaling

**DOI:** 10.1101/303990

**Authors:** Jacob C Garza, Xiaoli Qi, Klaudio Gjeluci, Melanie P Leussis, Himanish Basu, Surya A Reis, Wen Ning Zhao, Nicolas H Piguel, Peter Penzes, Stephen J Haggarty, Gerard J Martens, Geert Poelmans, Tracey L Petryshen

## Abstract

The ankyrin 3 gene (*ANK3*) is a well-established risk gene for psychiatric illness, but the mechanisms underlying its pathophysiology remain elusive. We examined the molecular effects of disrupting brain-specific *Ank3* isoforms in mouse and neuronal model systems. RNA sequencing of hippocampus from *Ank3+/-* and *Ank3+/+* mice identified altered expression of 282 genes that were enriched for microtubule-related functions. Results were supported by increased expression of microtubule end-binding protein 3 (EB3), an indicator of microtubule dynamics, in *Ank3+/-* mouse hippocampus. Live-cell imaging of EB3 movement in primary neurons from *Ank3+/-* mice revealed impaired elongation of microtubules. Using a CRISPR-dCas9-KRAB transcriptional repressor in mouse neuro-2a cells, we determined that repression of brain-specific *Ank3* increased EB3 expression, decreased tubulin acetylation, and increased the soluble:polymerized tubulin ratio, indicating enhanced microtubule dynamics. These changes were rescued by inhibition of glycogen synthase kinase 3 (GSK3) with lithium or CHIR99021, a highly selective GSK3 inhibitor. Brain-specific *Ank3* repression in neuro-2a cells increased GSK3 activity (reduced inhibitory phosphorylation) and elevated collapsin response mediator protein 2 (CRMP2) phosphorylation, a known GSK3 substrate and microtubule-binding protein. Pharmacological inhibition of CRMP2 activity attenuated the rescue of EB3 expression and tubulin polymerization in *Ank3* repressed cells by lithium or CHIR99021, suggesting microtubule instability induced by *Ank3* repression is dependent on CRMP2 activity. Taken together, our data indicate that aNK3 functions in neuronal microtubule dynamics through GSK3 and its downstream substrate CRMP2. These findings reveal cellular and molecular mechanisms underlying brain-specific ANK3 disruption that may be related to its role in psychiatric illness.

## Introduction

Large-scale genomic studies are providing a clearer picture of the genetic architecture of psychiatric illness. Genetic variation in *ANK3* is associated with several psychiatric disorders, including bipolar disorder (BD) and autism spectrum disorders (ASD).^1–11^ Human postmortem brain studies demonstrate that carriers of *ANK3* alleles associated with BD have lower *ANK3* expression at the transcript and protein levels,^12,13^ suggesting that decreased expression of aNK3 contributes to disease. Despite strong genetic evidence that *ANK3* contributes to psychiatric illness^14^, the precise mechanism is unknown.

*ANK3* encodes the ankyrin-G scaffolding protein that anchors integral membrane proteins to the cytoskeleton.^15,16^ There are several protein isoforms of ankyrin-G due to alternative splicing and alternative starting exons.^13,17^ These isoforms have unique functions and tissue distribution, including isoforms that are only expressed in brain. Genomic regions associated with BD span exon 1b of *ANK3* and the intron upstream of exon 37, exons that are only present in brain-specific isoforms. Furthermore, rare mutations identified in ASD patients are predominantly located within the brain-specific exon 37.^1–8^ The brain-specific isoforms are primarily known for their function in formation of the neuron axon initial segment (AIS) and clustering of ion channels at the nodes of Ranvier along axons.^18,19^ Interestingly, BD patients carrying a BD-associated risk allele for *ANK3* have a decreased fractional anisotropy in the uncinate fasciculus, which is an indication of impaired axon function or axonal damage in these forebrain connections.^11^

Recent evidence implicates cytoskeleton dysfunction in psychiatric illness.^20–22^ Microtubules are components of the cytoskeleton that contribute to the morphology of axons and dendrites in neurons, and facilitate transport of cellular cargo. They are composed of α and β tubulin heterodimers that continuously polymerize and depolymerize at the microtubule plus end (i.e., microtubule dynamics), leading to continuous growth and shrinkage of microtubules.^23^ ANK3 is reported to bind microtubules directly or through binding of microtubule associated proteins at the plus end stabilizing cap^24–26^ that prevents depolymerization. The interaction between ANK3 and microtubules provides a biological basis for examining brain-specific *Ank3* in the regulation of microtubule dynamics.

We have previously demonstrated that brain-specific *Ank3* isoforms regulate psychiatric-related behaviors in mice, and alterations in these behaviors in *Ank3+/-* mice are reversed by the mood stabilizer lithium.^27,28^ An important target of lithium is GSK3, which is implicated in psychiatric illness by animal studies.^29,30^ Among the downstream substrates of GSK3, CRMP2 has emerged as a prime target for regulation of microtubule dynamics and stability.^31–33^ In its unphosphorylated state, CRMP2 binds tubulin heterodimers and stabilizes the plus end of microtubules;^34^ however, upon phosphorylation by GSK3, CRMP2 activity is suppressed and binding to microtubules is reduced. In *Caenorhabditis elegans*, the homologs of *ANK3* (UNC-44) and *CRMP2* (UNC-33) are required to organize microtubules in neurons.^35^ Therefore, it is possible that ANK3 modulates microtubule dynamics through a mechanism involving CRMP2 signaling.

In the current study, we investigated the molecular impact of reducing expression of brain-specific *Ank3* isoforms, based on the patient genetic and expression studies noted above that implicate reduced expression of these isoforms in disease. Using RNA sequencing, biochemical, and live cell-imaging methods in mouse and neuronal model systems, we determined that brain-specific *Ank3* deficiency is associated with enhanced microtubule dynamics (i.e., increased tubulin polymerization/depolymerization). Furthermore, we demonstrate that microtubule changes induced by brain-specific *Ank3* repression are rescued by lithium or a selective inhibitor of GSK3 through a CRMP2-dependent mechanism. Our findings establish for the first time that brain-specific *Ank3* is important for maintaining proper microtubule dynamics through GSK3/CRMP2 signaling.

## Materials and Methods

See Supplementary Information for detailed methods.

### Animals

Male *Ank3+/-* mice with heterozygous disruption of *Ank3* exon 1b^18^ were crossed to female C57BL/6J mice (Jackson Laboratory, Bar Harbor, ME) to generate *Ank3+/-* and *Ank3+/+* progeny. Experiments were conducted in accordance with the National Institutes of Health guidelines and approval of the Institutional Animal Care and Use Committees of Massachusetts Institute of Technology and Massachusetts General Hospital.

### RNA sequencing and data analysis

Hippocampal RNA from 10 male 16-20wk old mice per genotype was pooled for RNA sequencing. Trimmed sequence reads were aligned onto the Mus musculus GRCm38/mm10 genome and analyzed using the Tuxedo package within the GenePattern platform (https://genepattern.broadinstitute.org).^36^ Differentially expressed genes were identified based on minimum 1.2-fold change and uncorrected *P* ≤ 1×10^−3^. The Ingenuity Pathway Analysis package was used to identify overrepresented biological pathways, with a focus on ‘Canonical Pathways’ and ‘Diseases and Functions’.

### Live cell imaging

Mouse primary forebrain neurons were generated from P0 *Ank3+/+* and *Ank3+/-* mice. Cells were transfected with mPA-GFP-EB3-7 (Addgene, Cambridge, MA) at DIV11-12 and imaged at DIV14. Axon segments 70-150 μm in length were imaged starting ~60 μm from the soma at 2 s intervals for 300 s. EB3 comet trajectory was manually traced from kymographs generated using the ImageJ Kymolyzer macro^37^ to calculate comet length, duration, and velocity.

### Western blot

Protein lysates from neuro-2a cells or hippocampal tissue from male mice were separated by SDS-PAGE and blotted onto PVDF membranes, probed with specific primary antibodies and HRP-linked secondary antibodies (Supplementary Table 1), followed by electrochemiluminescent detection. Protein expression was quantified by normalizing to GAPDH, and phosphorylated or acetylated protein expression by normalizing to the corresponding total protein.

### CRISPR-mediated *Ank3* transcriptional repression

Single guide RNA (sgRNA) sequences were designed to target mouse *Ank3* exon 1b using the CRISPR Design Tool (http://crispr.mit.edu) (Supplementary Table 2). *Ank3*-targeting and nontargeting control sgRNAs were cloned into the sgRNA(MS2)-EF1α plasmid (gift from Dr. Feng Zhang).^38^

### Cell culture

Mouse neuro-2a cells (ATCC, Manassas, VA) were dual transfected with the pHAGE-EF1α-dCas9-KRAB transcriptional repressor plasmid^39^ (Addgene) and sgRNA(MS2)-EF1α plasmid expressing *Ank3*-targeting or control sgRNA. For drug experiments, cells were treated for 1 h with the GSK3 inhibitors lithium (Sigma-Aldrich, St. Louis, Mo) or CHIR99021 (LC Laboratories, Woburn, Ma), or for 24 h with the CRMP2 inhibitor lacosamide (Sigma-Aldrich, St. Louis, Mo).

### RT-qPCR

SYBR Green qPCR was performed using 1 μg cDNA using gene and *Ank3* isoform-specific primers (Supplementary Table 3).^40^ Expression was normalized to beta-2-microglobulin.

### Tubulin polymerization assay

Neuro-2a cells were lysed to obtain soluble and insoluble protein fractions for Western blot detection of α-tubulin.^41,42^

### Statistical analysis

Statistical analyses were performed by unpaired Student’s *t*-test, or one- or two-way ANOVA followed by *post hoc* tests, using StatView version 5 or SPSS version 16. Sample sizes were determined based on previously published literature and our preliminary data. Significance threshold was set at *P*<0.05.

## Results

### RNA sequencing identifies expression changes in microtubule-related pathways in *Ank3+/-* mouse hippocampus

RNA sequencing analysis was performed to identify genes with altered expression in hippocampus from *Ank3+/-* mice, which exhibit 50% reduced expression of brain-specific *Ank3* compared to wild-type *Ank3+/+* mice (Supplementary Figure 1). Based on our targeted read depth of 25 million reads, we expected the RNA sequencing analysis to detect predominantly abundant genes. A total of 282 genes were significantly differentially expressed (fold change ≥1.2, uncorrected *P* ≤ 1×10^−3^) between *Ank3+/-* and *Ank3+/+* hippocampus (102 upregulated, 180 downregulated; Supplementary Table 4). Ingenuity Pathway Analysis of the differentially expressed genes identified significant overrepresentation of the Axonal Guidance Signaling canonical pathway (17 genes; corrected *P*=0.0088). Among the Disease & Function pathways, the Microtubule Dynamics pathway was most significantly enriched (34 genes; uncorrected *P*=6.7×10^−5^). Ten of the 17 Axonal Guidance Signaling genes were also annotated as Microtubule Dynamics genes. The Microtubule Dynamics pathway is categorized within a higher-level function of Cellular Assembly and Organization, which contains three other pathways that were overrepresented among the 282 differentially expressed genes, although with weaker statistical evidence: Fusion of Vesicles (6 genes, *P*=2.0×10^−4^), Organization of Cytoskeleton (36 genes, *P*=3.1×10^−4^), and Extension of Cellular Protrusions (10 genes, *P*=3.3×10^−4^). Many of the genes in the latter three pathways overlap the Microtubule Dynamics and Axonal Guidance Signaling pathways, resulting in a total of 47 genes across the five pathways (Table 1). As the identified pathways represent cellular functions requiring microtubules, we focused subsequent experiments on the role of *Ank3* in microtubule dynamics.

**Table 1.**
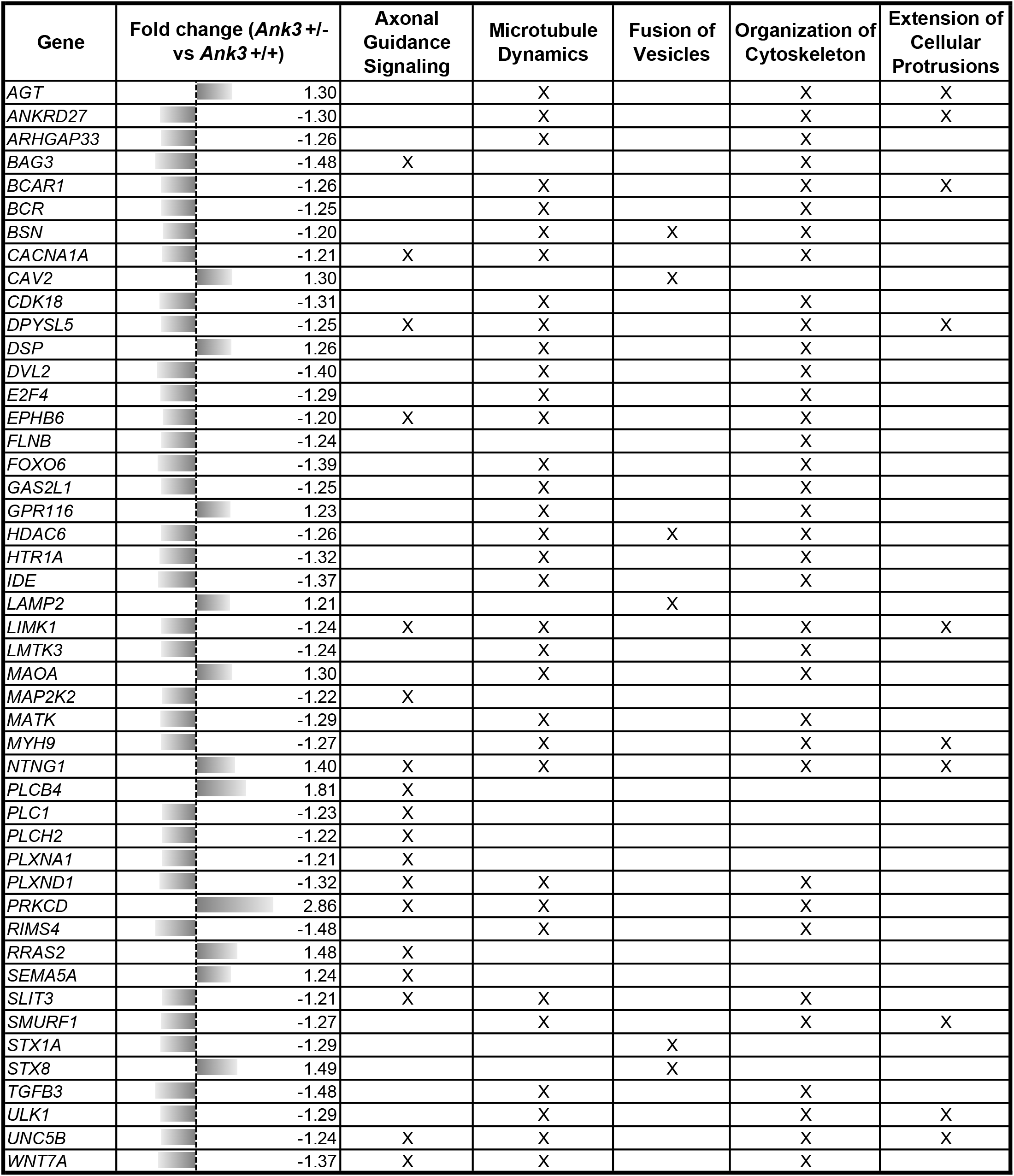
Microtubule-related pathways enriched for genes that are differentially expressed between *Ank3+/-* and *Ank3+/+* mouse hippocampus.

| Gene | Fold change (*Ank3 +/-* vs *Ank3 +/+*) | Axonal Guidance Signaling | Microtubule Dynamics | Fusion of Vesicles | Organization of Cytoskeleton | Extension of Cellular Protrusions |
| --- | --- | --- | --- | --- | --- | --- |
| AGT | 1.30 |  | X |  | X | X |
| ANKRD27 | -1.30 |  | X |  | X | X |
| ARHGAP33 | -1.26 |  | X |  | X |  |
| BAG3 | -1.48 | X |  |  | X |  |
| BCAR1 | -1.26 |  | X |  | X | X |
| BCR | -1.25 |  | X |  | X |  |
| BSN | -1.20 |  | X | X | X |  |
| CACNA1A | -1.21 | X | X |  | X |  |
| CAV2 | 1.30 |  |  | X |  |  |
| CDK18 | -1.31 |  | X |  | X |  |
| DPYSL5 | -1.25 | X | X |  | X | X |
| DSP | 1.26 |  | X |  | X |  |
| DVL2 | -1.40 |  | X |  | X |  |
| E2F4 | -1.29 |  | X |  | X |  |
| EPHB6 | -1.20 | X | X |  | X |  |
| FLNB | -1.24 |  |  |  | X |  |
| FOXO6 | -1.39 |  | X |  | X |  |
| GAS2L1 | -1.25 |  | X |  | X |  |
| GPR116 | 1.23 |  | X |  | X |  |
| HDAC6 | -1.26 |  | X | X | X |  |
| HTR1A | -1.32 |  | X |  | X |  |
| IDE | -1.37 |  | X |  | X |  |
| LAMP2 | 1.21 |  |  | X |  |  |
| LIMK1 | -1.24 | X | X |  | X | X |
| LMTK3 | -1.24 |  | X |  | X |  |
| MAOA | 1.30 |  | X |  | X |  |
| MAP2K2 | -1.22 | X |  |  |  |  |
| MATK | -1.29 |  | X |  | X |  |
| MYH9 | -1.27 |  | X |  | X | X |
| NTNG1 | 1.40 | X | X |  | X | X |
| PLCB4 | 1.81 | X |  |  |  |  |
| PLC1 | -1.23 | X |  |  |  |  |
| PLCH2 | -1.22 | X |  |  |  |  |
| PLXNA1 | -1.21 | X |  |  |  |  |
| PLXND1 | -1.32 | X | X |  | X |  |
| PRKCD | 2.86 | X | X |  | X |  |
| RIMS4 | -1.48 |  | X |  | X |  |
| RRAS2 | 1.48 | X |  |  |  |  |
| SEMA5A | 1.24 | X |  |  |  |  |
| SLIT3 | -1.21 | X | X |  | X |  |
| SMURF1 | -1.27 |  | X |  | X | X |
| STX1A | -1.29 |  |  | X |  |  |
| STX8 | 1.49 |  |  | X |  |  |
| TGFB3 | -1.48 |  | X |  | X |  |
| ULK1 | -1.29 |  | X |  | X | X |
| UNC5B | -1.24 | X | X |  | X | X |
| WNT7A | -1.37 | X | X |  | X |  |

### Enhanced microtubule dynamics in *Ank3+/-* mouse hippocampus

To obtain support for microtubule defects in *Ank3+/-* mouse brain, we evaluated the expression of the EB3 end-binding protein. EB3 modulates stability at the microtubule plus end, serving as a marker of growing microtubules,^43^ and is reported to directly interact with Ank3.^24–26^ Western blot analysis of EB3 in hippocampus isolated from *Ank3+/+* and *Ank3+/-* mice determined that expression was increased 1.5-fold in *Ank3+/-* mice compared to *Ank3+/+* mice (*P*<0.001, Figure 1). The substantial elevation in EB3 expression suggests that reduction of brain-specific *Ank3* is associated with enhanced microtubule dynamics (i.e., increased polymerization/depolymerization of tubulin at the plus end).

**Figure 1.**
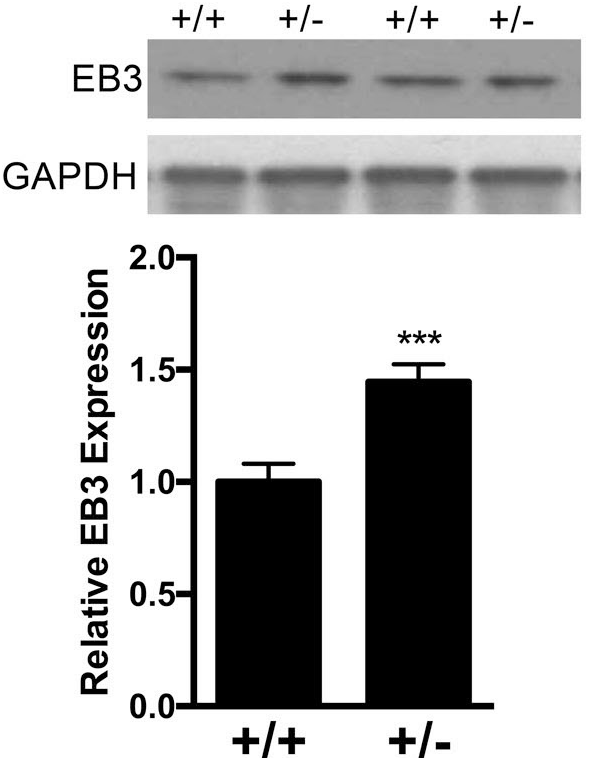
Increased expression of EB3 in *Ank3+/-* mouse hippocampus. Top: Representative Western blot of EB3 and GAPDH reference protein in hippocampal tissue from *Ank3+/+* and *Ank3+/-* mice. Bottom: Quantification of EB3 expression normalized to GAPDH expression. Data were analyzed using two-tailed Student’s *t*-test, *Ank3+/+* n=12; *Ank3+/-* n=13. Data are presented as mean ± s.e.m. \*\*\**P*<0.001.

### Brain-specific *Ank3* reduction impairs microtubule elongation in axons

To directly monitor the effect of *Ank3* reduction on the dynamic properties of microtubules, we performed live-cell imaging of GFP-tagged EB3 in mouse primary neurons. As microtubules polymerize, EB3-GFP puncta at the plus end cap appear as mobile comets, which dissipate when the microtubules depolymerize.^44^ We analyzed EB3 comet length, duration, and velocity in axons of DIV14 primary neurons from *Ank3+/+* and *Ank3+/-* neonatal mice (Figure 2a). The trajectory length of EB3 comets was ~15% shorter in axons of *Ank3+/-* neurons compared to *Ank3+/+* neurons (Figure 2b, *P*=0.002), indicating decreased microtubule elongation. EB3 comets in *Ank3+/-* axons were detected in ~15% fewer frames of the 300s time-lapse imaging video compared to the comets in *Ank3+/+* axons (Figure 2c, *P*=0.003), suggesting decreased duration of EB3 bound to microtubule plus ends. The velocity of EB3 comets did not differ between *Ank3+/-* axons and *Ank3+/+* axons, indicating that the rate of microtubule polymerization was unchanged (Figure 2d, *P*>0.05). These data suggest an increase in the dynamics of microtubules with an overall reduction of microtubule elongation in *Ank3+/-* axons compared to *Ank3+/+* axons.

**Figure 2.**
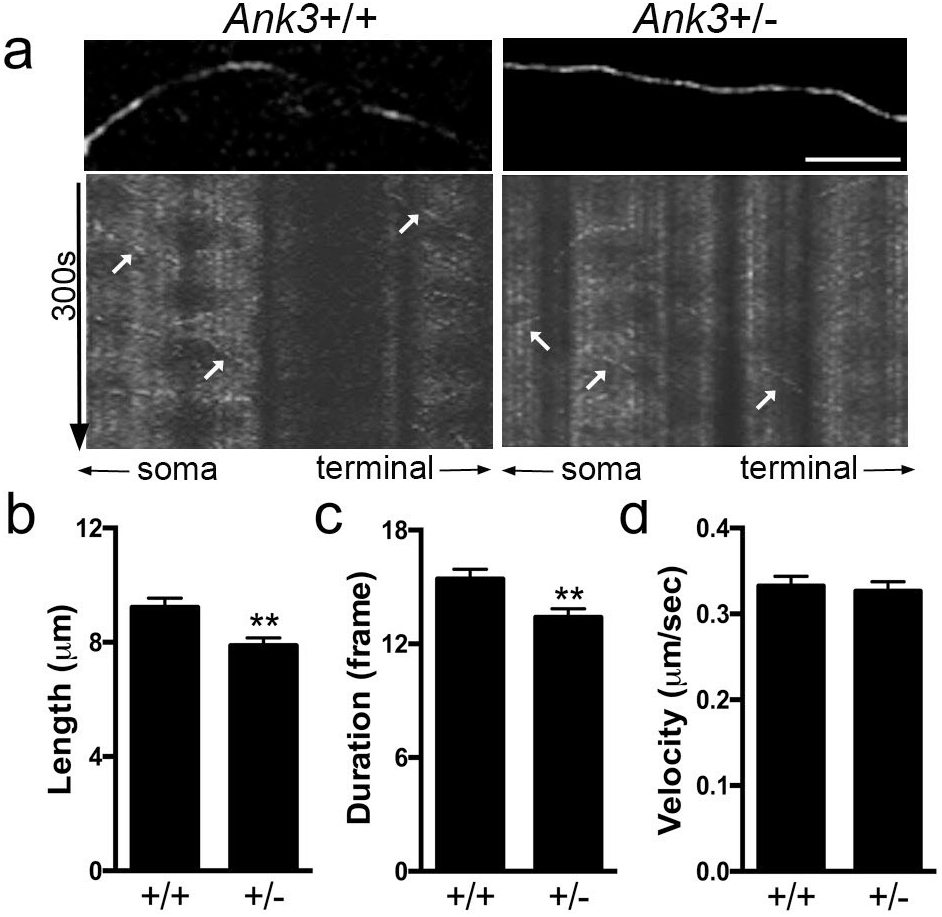
Reduced brain-specific *Ank3* expression is associated with enhanced microtubule dynamics in axons of mouse primary neurons, **(a)** Representative kymographs from live-cell imaging of EB3 comets in DIV14 primary neurons from *Ank3+/+* (left) and *Ank3+/-* (right) P0 neonatal mice. Top: EB3-GFP puncta present in the axon. Scale bar = 10 μm. Bottom: Kymographs show the movement of EB3 comets. The x axis represents the position along the axon and the y axis represents time (300s). White arrows indicate representative comet traces. Quantification of EB3 comet **(b)** length, **(c)** duration, and **(d)** velocity. Data were collected from three independent experiments. *Ank3+/+, n*=245 comets from 27 axons of 6 mice; *Ank3+/-, n*=220 comets from 14 axons of 7 mice. Data were analyzed using two-tailed Student’s *t*-test. Data are presented as mean ± s.e.m. \*\**P*<0.01.

### Establishment of a neuronal model of brain-specific *Ank3* repression

We established a cellular model to investigate the mechanism underlying impaired microtubule elongation associated with brain-specific *Ank3* reduction. Mouse neuro-2a cells were dualtransfected with a CRISPRIdCas9 KRAB repressor and either a sgRNA targeting *Ank3* exon 1b or a control sgRNA (Figure 3a). Fourteen sgRNA sequences were screened for efficacy of *Ank3* exon1b repression (Supplementary Figure 2). One sgRNA (#7) was selected and selective repression of Ank3 exon 1b was verified (Figure 3b; Supplementary Information). Western blot analysis indicated that *Ank3* repression elevated EB3 expression by 40% compared to control cells (Figure 3c, *P*<0.01). This aligns with our earlier observation of increased EB3 expression in hippocampus from *Ank3+/-* mice compared to *Ank3+/+* mice (Figure 1), thereby validating our neuronal model system.

**Figure 3.**
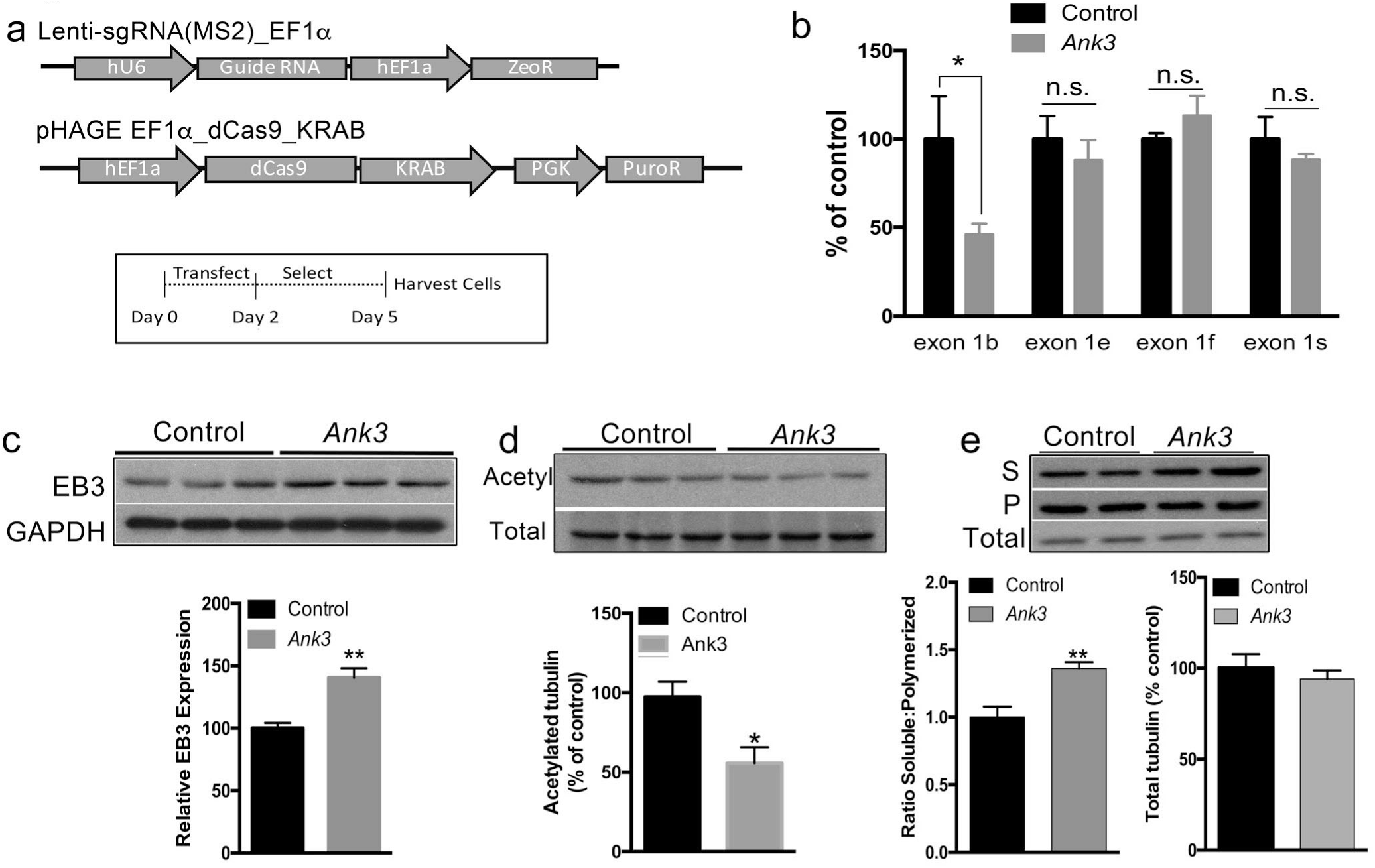
CRISPR/dCas9 mediated repression of brain-specific *Ank3* in mouse neuro-2a cells, **(a)** Top: Schematic representation of the plasmids used for repression of brain-specific *Ank3*. pHAGE-EF1α_dCas9_KRAB expressed the deactivated Cas9 fused to the KRAB transcriptional repressor. Lenti-sgRNA(MS2)_EF1α expressed the control sgRNA or an sgRNA targeting *Ank3* exon 1b. See Supplementary Figure 2 for qPCR screening of sgRNA sequences for efficiency of *Ank3* exon 1b repression. Bottom: Schematic representation of the experimental design. Neuro-2a cells were transfected with the two plasmids, followed by puromycin and zeocin selection, and cell harvest for protein extraction, **(b)** Expression of *Ank3* exon 1b, exon le, exon 1f, and exon 1s was measured by qPCR using isoform-specific primers in neuro-2a cells expressing the control sgRNA or the *Ank3*-targeting sgRNA. Expression was normalized to beta-2-microglobulin and presented as percent of the control sgRNA for each starting exon, **(c)** Transcriptional repression of brain-specific *Ank3* in neuro-2a cells alters EB3 expression. Top: Representative Western blots of EB3 and GAPDH. Bottom: Quantification of EB3 expression normalized to GAPDH. **(d)** Transcriptional repression of brain-specific *Ank3* in neuro-2a cells alters acetylation of α-tubulin. Top: Representative Western blots of acetylated α-tubulin (Acetyl) and total α-tubulin (Total). Bottom: Quantification of acetylated α-tubulin expression normalized to total tubulin, **(e)** Transcriptional repression of brain-specific *Ank3* in neuro-2a cells alters the ratio of soluble:polymerized tubulin. Top: Representative Western blots of α-tubulin in soluble (S) and polymerized (P) protein fractions, and in total cell lysate (Total). Bottom left: Quantification of the ratio of soluble:polymerized tubulin. Bottom right: Quantification of total tubulin normalized to GAPDH. Western blot data were averaged from three independent experiments with three biological replicates per group in each experiment. Control, non-targeting sgRNA; *Ank3*, sgRNA targeting *Ank3* exon 1b. Data were analyzed using two-tailed Student’s *t*-test. Data are presented as mean ± s.e.m. \**P*<0.05, ** *P*<0.01, n.s. indicates not significant.

### Repression of brain-specific *Ank3* in cells reduces polymerized tubulin

To assess the enhancement of microtubule dynamics associated with brain-specific *Ank3* repression, we examined characteristics of tubulin (i.e. acetylation and polymerization state) in our neuronal model system. Tubulin acetylation is an indicator of the overall stability of microtubules, such that microtubules that are more stable and resistant to turnover have higher acetylation levels, whereas more microtubules that are more dynamic and susceptible to turnover have lower acetylation levels.^45,46^ Western blot analysis revealed a 45% reduction of acetylated tubulin normalized to total tubulin in *Ank3* repressed cells compared to control cells (Figure 3d, *P*<0.05), suggesting microtubules are more dynamic when *Ank3* is repressed. Further, Western blot analysis of tubulin in soluble and polymerized protein fractions^41,47–49^ determined that brain-specific *Ank3* repression led to an increase in the amount of tubulin in the soluble protein fraction and a concomitant decrease in the polymerized fraction compared to control cells. This resulted in a 1.4-fold increase in the ratio of soluble to polymerized tubulin (Figure 3e, *P*<0.01), indicating a shift in equilibrium from microtubule-associated tubulin towards free tubulin as a result of brain-specific *Ank3* repression. This shift was not due to changes in total tubulin expression between *Ank3*-repressed and control cells (Figure 3e, *P*>0.05). These results are consistent with increased turnover at the microtubule plus ends due to repression of brain-specific *Ank3*.

### GSK3 inhibition rescues microtubule changes associated with *Ank3* repression

To examine the molecular mechanisms underlying enhanced microtubule dynamics in our neuronal model system, we investigated the effects of lithium treatment, which we previously demonstrated reverses behavioral abnormalities of brain-specific *Ank3+/-* mice.^27,28^ In addition, to determine whether any effects of lithium are mediated by its known target GSK3, we also investigated CHIR99021, a highly selective inhibitor of both GSK3α and GSK3β.^50^ Control and *Ank3* exon 1b repressed neuro-2a cells treated with vehicle, 1mM lithium chloride, or 1μM CHIR99021 for 1h (Figure 4a) were assessed for EB3 expression and tubulin polymerization. While *Ank3* repression increased EB3 expression by ~50% compared to control in vehicle-treated cells (Figure 4b, *post hoc P*<0.01), as expected based on our previous experiment (Figure 3e), the increase was attenuated by lithium and CHIR99021 treatment (Figure 4b, both *post hoc P*>0.05). Similarly, while *Ank3* exon 1b repression increased the ratio of soluble:polymerized tubulin by 1.35 fold compared to control in vehicle-treated cells (Figure 4c, *post hoc P*<0.01), the ratio was normalized by treatment with lithium or CHIR99021 (Figure 4c, both *post hoc P*>0.05). To rule out the possibility that the rescue was due to normalizing expression of *Ank3*, we confirmed that *Ank3* expression did not differ after treatment with lithium or CHIR99021 compared to vehicle (Supplementary Figure 3). These results indicate that GSK3 is involved in changes to microtubule dynamics induced by repression of brain-specific *Ank3*.

**Figure 4.**
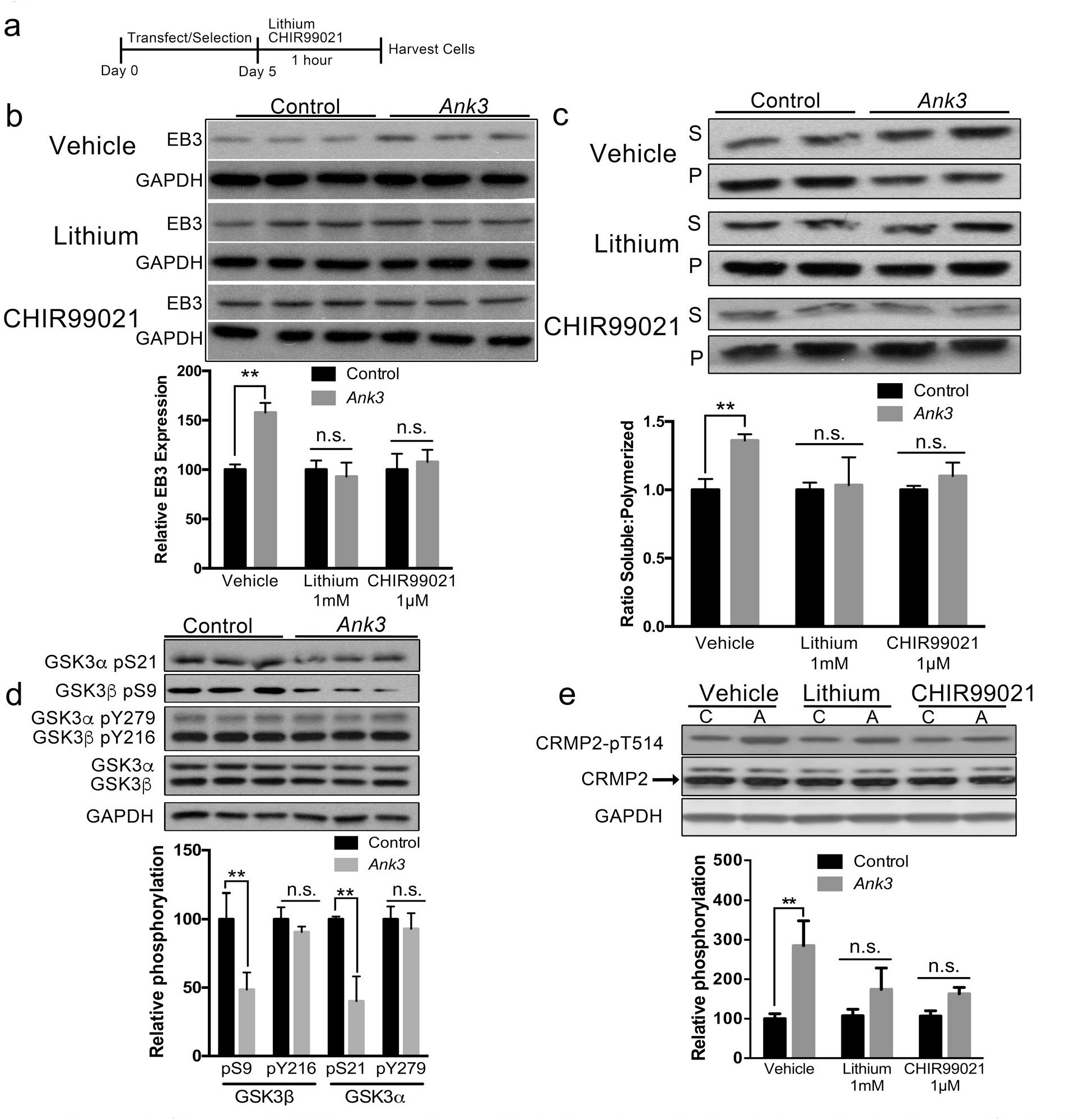
Changes in EB3 expression and tubulin polymerization induced by brain-specific *Ank3* repression are normalized by inhibition of GSK3. **(a)** Schematic representation of the experimental design. Mouse neuro-2a cells were transfected with pHAGE-EF1α-dCas9-KRAB repressor plasmid and sgRNA(MS2)_EF1α plasmid expressing either the non-targeting control sgRNA or the sgRNA targeting *Ank3* exon 1b, followed by puromycin and zeocin selection, and treatment with lithium (1mM), CHIR99021 (1μM), or DMSO vehicle for 1hr prior to cell harvest for protein extraction, **(b)** Top: Representative Western blot of EB3 and GAPDH. Bottom: Quantification of EB3 expression normalized to GAPDH. Two-way ANOVA, drug effect F_(2,30)_=4.217 *P*=0.02, *Ank3* repression effect F_(1,30)_=4.137 *P*=0.05, interaction F_(2,30)_=4.217, *P*=0.02. **(c)** Top: Representative Western blot of α-tubulin in soluble (S) and polymerized (P) protein fractions. Bottom: Quantification expressed as the ratio of α-tubulin in soluble and polymerized fractions. Two-way ANOVA, drug effect F_(2,28)_=2.460 *P*=0.10, *Ank3* repression effect F_(1,28)_=3.281 *P*=0.08, interaction F_(2,28)_=5.463 *P*=0.01. **(d)** Top: Representative Western blot of GSK3α/β phosphorylation at serine 21 (GSK3α-pS21), serine 9 (GSK3β-pS9), tyrosine 279 (GSK3α-pY279), and tyrosine 216 (GSK3β-pY216), total GSK3α and GSK3β, and GAPDH. Bottom: Quantification of GSK3β-pS9 and GSK3β-pY216 normalized to total GSK3β, and GSK3α-pS21 and GSK3α-pY279 normalized to total GSK3α. **(e)** Top: Representative Western blot of CRMP2 phosphorylation at threonine 514 (CRMP2-pT514), total CRMP2, and GAPDH. Bottom: Quantification of CRMP2-pT514 normalized to total CRMP2. Two-way ANOVA, drug effect F_(2,12)_=4.137 *P*=0.05, *Ank3* repression effect F_(1,12)_=3.281 *P*=0.08, interaction F_(2,12)_=4.217 *P*=0.02. Western blot data were averaged from three independent experiments with three biological replicates per group in each experiment. Control or C, non-targeting sgRNA; *Ank3* or A, sgRNA targeting *Ank3* exon 1b. Data were analyzed using two-tailed Student’s *t*-test or ANOVA and Bonferroni *post hoc* tests. Data are presented as mean ± s.e.m. * *P*<0.05, \*\**P*<0.01. n.s. indicates not significant.

To investigate the relationship between brain-specific *Ank3* and GSK3 activity, we measured phosphorylation at key regulatory sites of GSK3β and GSK3α, serine 9 (GSK3β-pS9) and serine 21 (GSK3α-pS21), which suppress activity,^51,52^ and tyrosine 216 (GSK3β-pY216) and tyrosine 279 (GSK3α-pY279), which promote activity in the absence of serine 9/21 phosphorylation.^53^ While repression of brain-specific *Ank3* did not affect GSK3β-pY216 or GSK3α-pY279 levels (Figure 4d, both *P*>0.05), there were significant reductions in GSK3β-pS9 and GSK3α-pS21 (Figure 4d, both *P*<0.01). These data suggest that repression of brain-specific *Ank3* upregulates GSK3 activity through reduced phosphorylation of GSK3β-pS9 and GSK3α-pS21 regulatory sites.

### Brain-specific *Ank3* repression enhances GSK3-mediated inhibition of CRMP2 microtubule stabilization

The GSK3 substrate CRMP2 has a key regulatory role in microtubule dynamics by promoting microtubule stability through interactions with tubulin heterodimers and acting as an adapter with motor proteins.^33,54^ This interaction is highly regulated by GSK3-dependent phosphorylation, where phosphorylation of CRMP2 threonine 514 (CRMP2-pT514) inhibits the interaction of CRMP2 with tubulin heterodimers.^33,55^ We investigated the effect of brain-specific *Ank3* repression on CRMP2-pT514 and whether it was modified by inhibition of GSK3 in our neuronal model system. While *Ank3* exon 1b repression in neuro-2a cells resulted in no change in total CRMP2 levels compared to control cells (*P*>0.05), inhibitory CRMP2-pT514 was significantly increased nearly 3-fold (Figure 4e; *post hoc* vehicle-control vs vehicle-repressor *P*<0.01). As expected, treatment with lithium (1mM) or CHIR99021 (1μM) attenuated the increase in CRMP2-pT514 induced by *Ank3* exon 1b repression, such that there was no significant difference between the *Ank3* repressed and control groups (Figure 4e, *post hoc* lithium-control vs lithium-repressor *P*>0.05, *post hoc* CHIR99021-control vs CHIR99021 - repressor *P*>0.05). These results suggest that repression of brain-specific *Ank3* increases EB3 expression and soluble:polymerized tubulin ratio through GSK3-mediated inhibition of CRMP2.

### Rescue of enhanced microtubule dynamics induced by *Ank3* repression requires CRMP2 activity

To address whether CRMP2 activity is required for GSK3 inhibition to rescue microtubule changes induced by *Ank3* repression, we evaluated whether lacosamide blocks the rescue by lithium or CHIR99021 in our neuronal model system. Lacosamide affects slow activation of sodium channels at doses of 100-500μM, but lower doses of 3-5μM inhibit CRMP2 activity and tubulin binding without affecting sodium channel activation.^56–59^ Pre-treatment with a low 5μM dose of lacosamide 24hr prior to 1mM lithium or 1μM CHIR99021 treatment (Figure 5a) blocked lithium and CHIR99021 from rescuing changes in EB3 (Figure 5b) and steady state tubulin polymerization (Figure 5c) induced by *Ank3* exon 1b repression. Specifically, when EB3 expression was evaluated by Western blot, ANOVA and *post hoc* analysis revealed that, in the vehicle treated groups, *Ank3* repression compared to control increased EB3 expression (Figure 5b, vehicle-vehicle control vs vehicle-vehicle repressor *P*<0.01), which was rescued by treatment with lithium or CHIR99021 (Figure 5b, vehicle-lithium control vs vehicle-lithium repressor *P*>0.05, vehicle-CHIR99021 control vs vehicle-CHIR99021 repressor *P*>0.05). In contrast, in the lacosamide pre-treated groups, *Ank3* repression increased EB3 compared to control (Figure 5b, lacosamide-vehicle control vs lacosamide-vehicle repressor *P*<0.05), but the increase was not rescued by lithium or CHIR99021 (Figure 5b, lacosamide-lithium control vs lacosamide-lithium repressor *P*<0.05, lacosamide-CHIR99021 control vs lacosamide-CHIR99021 repressor *P*<0.01). Similarly, when the ratio of soluble:polymerized tubulin was assessed, ANOVA and *post hoc* analysis determined that, in the vehicle treated groups, Ank3 repression increased the ratio of soluble:polymerized tubulin (Figure 5c, vehicle-vehicle control vs vehicle-vehicle repressor *P*<0.05), which was rescued by lithium or CHIR99021 (Figure 5c, vehicle-lithium control vs vehicle-lithium repressor *P*>0.05, vehicle-CHIR99021 control vs vehicle-CHIR99021 repressor *P*>0.05). In contrast, in the lacosamide pre-treated groups, *Ank3* repression increased the ratio of soluble:polymerized tubulin (Figure 5c, lacosamide-vehicle control vs lacosamide-vehicle repressor *P*<0.05), but the increase was not rescued by lithium or CHIR99021 (Figure 5c, lacosamide-lithium control vs lacosamide-vehicle repressor *P*<0.01, lacosamide-CHIR99021 control vs lacosamide-CHIR99021 repressor *P*<0.01). Together, these data indicate that CRMP2 activity is required for GSK3 inhibition to rescue enhanced microtubule dynamics associated with repression of brain-specific *Ank3*.

**Figure 5.**
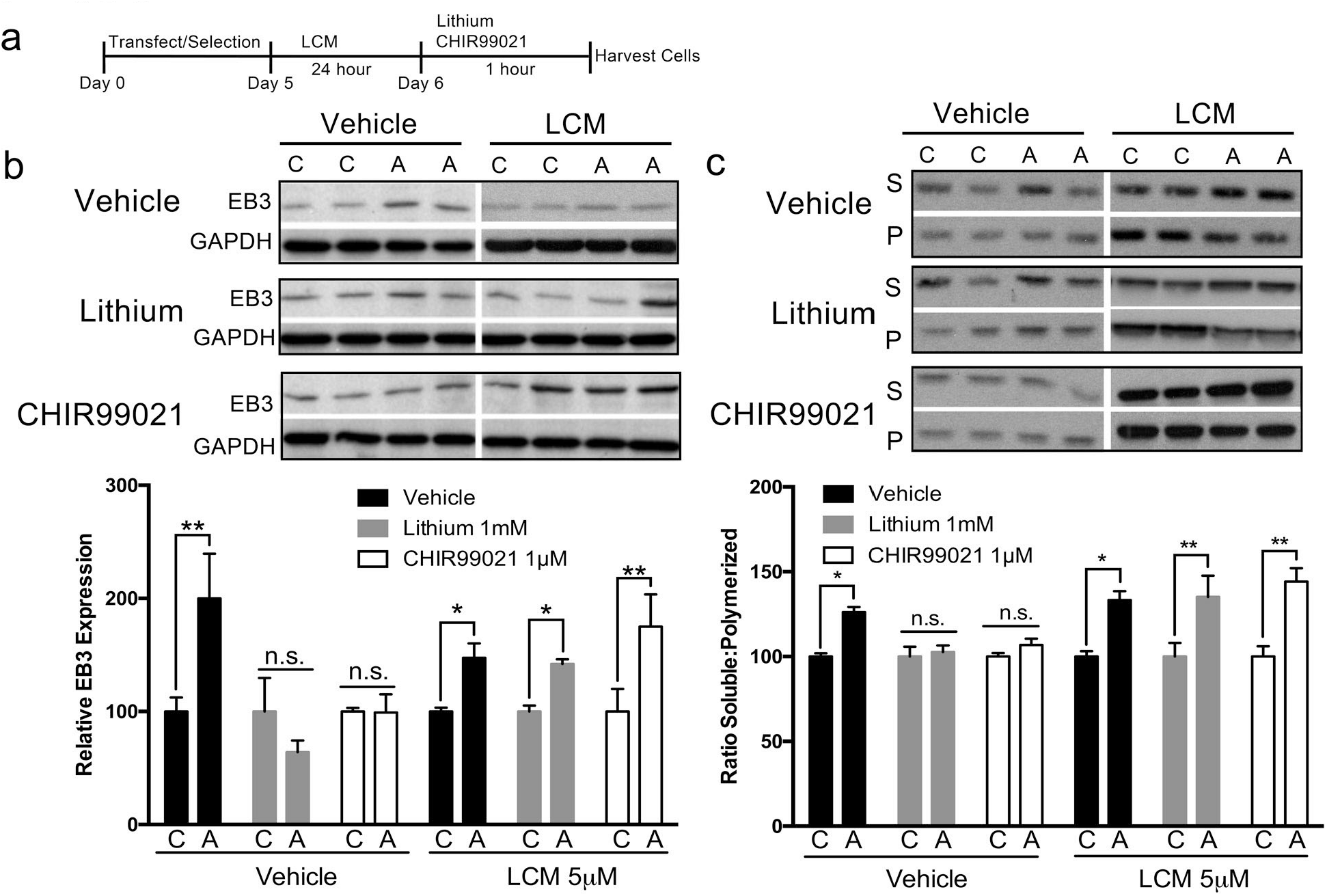
Rescue of altered EB3 expression and tubulin polymerization induced by brain-specific *Ank3* repression is blocked by inhibition of CRMP2. **(a)** Schematic representation of the experimental design. Mouse neuro-2a cells were transfected with pHAGE-EF1α-dCas9-KRAB repressor plasmid and sgRNA(MS2)_EF1α plasmid expressing either the non-targeting control sgRNA or the sgRNA targeting *Ank3* exon 1 b, followed by puromycin and zeocin selection. Cells were subsequently treated with 5μM lacosamide (LCM) for 24h, followed by treatment with lithium (1mM) or CHIR99021 (1μM) for 1h, and cell harvest for protein extraction, **(b)** Top: Representative Western blot of EB3 and GAPDH. Bottom: Quantification of EB3 expression normalized to GAPDH. Univariate ANOVA, LCM effect F_(1,60)_=2.00 *P*=0.163, lithium/CHIR99021 effect F_(5,60)_=3.586 *P*=0.03, *Ank3* repression effect F_(1,60)_=21.389 *P*<0.001, LCM and lithium/CHIR99021 interaction F_(2,60)_=7.505 *P*=0.001. **(c)** Top: Representative Western blot of α-tubulin in soluble (S) and polymerized (P) protein fractions. Bottom: Quantification of the ratio of soluble:polymerized tubulin. Univariate ANOVA, LCM effect F_(1,60)_=17.561 *P*<0.001, lithium/CHIR99021 effect F_(5,60)_=0.203 *P*=0.817, *Ank3* repression effect F_(1,60)_=68.479 *P*<0.001, LCM and lithium/CHIR99021 interaction F_(2,60)_=6.113 *P*=0.004. Western blot data were averaged from two independent experiments with three biological replicates per group in each experiment. C, control sgRNA; A, *Ank3*-targeting sgRNA. Data were analyzed using two-tailed Student’s *t*-test or ANOVA and Bonferroni *post hoc* tests. Data are presented as mean ± s.e.m. \**P*<0.05, \*\**P*<0.01. n.s. indicates not significant.

## Discussion

The current study used a multifaceted approach to identify and characterize the molecular functions of brain-specific *Ank3*. The key finding was that brain-specific *Ank3* is associated with microtubule dynamics via a GSK3/CRMP2-dependent mechanism (Figure 6). Our transcriptome-wide RNAseq analysis of *Ank3+/-* mouse hippocampus identified significantly altered expression of genes involved in pathways related to microtubule regulation and function, specifically axonal guidance signaling, microtubule dynamics, vesicle fusion, cytoskeletal organization, and extension of cellular protrusions. Subsequent live-cell imaging of primary neuron axons determined that microtubule dynamics are altered (i.e. decreased EB3 comet length and duration) in *Ank3+/-* mice compared to *Ank3+/+* mice. To examine the underlying molecular and biochemical basis, we utilized a CRISPR/dCas9-based neuronal model of brain-specific *Ank3* repression that exhibited changes in microtubule characteristics (increased EB3, increased ratio of soluble:polymerized tubulin, and decreased tubulin acetylation). The microtubule changes were rescued by inhibition of GSK3 and required active CRMP2, a GSK3 substrate that functions in microtubule stabilization. Taken together, this is the first study to demonstrate that brain-specific *Ank3* has a vital role in microtubule dynamics via a GSK3/CRMP2-dependent mechanism. Although it is not known whether *ANK3* contributes to psychiatric illness by altering microtubule function, it is intriguing that microtubules and microtubule regulators have previously been implicated in psychiatric disorders.^20^ Notably, microtubules are shortened and microtubule organization is perturbed in neuronal precursor cells from BD and schizophrenic patients, respectively.^60^ It will be important to investigate the relationship between *ANK3*, microtubules, and psychiatric illness in future studies.

**Figure 6.**
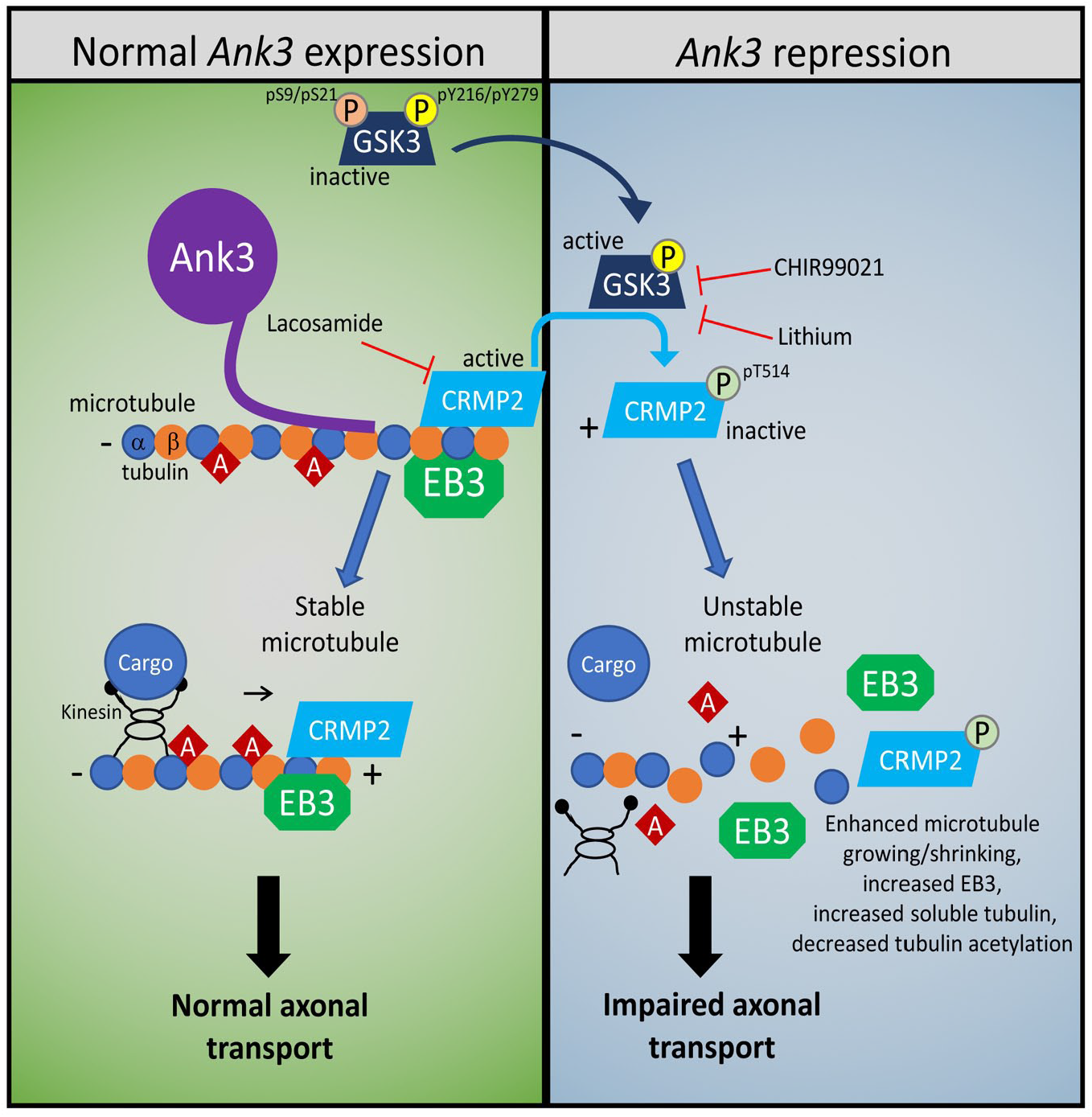
Brain-specific *Ank3* repression is associated with enhanced microtubule dynamics. Left: Under normal *Ank3* expression, microtubules are stabilized by binding of microtubule-associated proteins (MAPs), such as EB3 and CRMP2, to the microtubule plus end where α- and β-tubulin heterodimers polymerize to facilitate elongation of the microtubule. As microtubules elongate, EB3 and CRMP2 move along the growing plus end tip to stabilize newly generated microtubule segments. Acetylation (red diamond) accumulates on α-tubulin within the microtubule due to low microtubule turnover. Motor proteins, such as kinesin, mediate transport of cellular cargo along the microtubule towards the plus end, which is oriented towards the distal axon. Right: Repression of brain-specific *Ank3* reduces phosphorylation of GSK3 (pS9/pS21) and increases GSK3 activity, leading to an increase in CRMP2 phosphorylation (pT514) and impaired CRMP2 binding and stabilization of microtubules. Microtubules become more susceptible to catastrophes, as demonstrated by increased EB3 expression, reduced EB3 comet length and duration, and increased ratio of soluble:polymerized tubulin. The increased susceptibility to catastrophes increases microtubule turnover and decreases acetylation of α-tubulin. Inhibition of GSK3 activity by lithium or CHIR99021 reduces CRMP2 phosphorylation (pT514), thereby allowing CRMP2 to bind and stabilize microtubules. Pharmacological inhibition of CRMP2 by lacosamide reduces CRMP2 binding and stabilization of microtubules, which increases the amount of free tubulin and decreases acetylation of α-tubulin. Enhanced microtubule dynamics induced by brain-specific Ank3 repression may have a range of effects (e.g. axonal transport of synaptic vesicles, microtubule interaction with motor proteins) that alter neuronal function.

Live-cell imaging analyses determined that EB3 comet speed was unchanged in *Ank3+/-* mouse primary neuron axons, indicating that the rate of tubulin polymerization is normal. This suggests that tubulin heterodimers are able to form at microtubule plus ends when brain-specific *Ank3* is reduced. However, the observed decrease in length and duration of EB3 comets in *Ank3+/-* mouse primary neurons suggests impaired stability of growing microtubules. This is characteristic of increased microtubule catastrophes,^61,62^ i.e. switching from growth to rapid shortening, whereby the plus end stabilizing cap, where EB3 binds, is lost.^63^ The shift in tubulin equilibrium to a soluble state induced by *Ank3* repression, as observed in our neuronal model system, provides additional evidence for increased catastrophes, since diminished microtubule elongation would lead to accumulation of tubulin in the soluble pool rather than incorporation into growing microtubules. In support of this, treatment with taxol to stabilize microtubules has been reported to reduce the pool of soluble tubulin.^41,64^ Conversely, taxol-resistant cancer cells have a lower proportion of polymerized tubulin, as well as increased microtubule dynamics as indicated by increased movement of EB3 comets.^64^ Furthermore, additional data from our neuronal model shows diminished tubulin acetylation after *Ank3* repression, suggesting that microtubules are more susceptible to rapid turnover. These studies support our findings that reduced expression of brain-specific *Ank3* isoforms leads to increased activity at microtubule plus ends and instability of growing microtubules.

Among the lines of evidence implicating brain-specific *Ank3* in regulation of microtubule dynamics, we found that reduction of brain-specific *Ank3* in mice and our neuronal model system increased the overall expression of the EB3 microtubule end-binding protein. This is consistent with a recent report that *Ank3* knockdown in mouse primary hippocampal neurons increased EB3 puncta in axons.^65^ However, that study reduced expression of all *Ank3* isoforms by ~90%, whereas we targeted brain-specific *Ank3* isoforms that are specifically implicated in psychiatric illness,^1–8^ and we reduced expression by only ~50%, which is more consistent with patient brain expression changes.^12^ While the precise mechanism by which EB3 expression is increased following brain-specific *Ank3* repression is not known, several studies have shown that EB3 is elevated in response to enhanced microtubule dynamics,^66,67^ providing support that brain-specific *Ank3* repression is related to enhanced dynamics.

Our data suggest that *Ank3* reduction changes GSK3 and CRMP2 activity to modulate microtubule stability. Specifically, we found that repression of brain-specific *Ank3* resulted in a reduction of GSK3β-pS9 and GSK3α-pS21 (i.e. increased activity), and a concomitant increase in CRMP2-pT514 (i.e. decreased activity). This is interesting given our previous study demonstrating that lithium, which inhibits GSK3 in part through increasing pS9IpS21 levels through AKT activation,^50^ reversed psychiatric-like behaviors in *Ank3+/-* mice.^27^ GSK3 has been previously implicated in psychiatric illness^68–70^ however, further studies are warranted to elucidate how GSK3 activity, *Ank3*, and microtubules interact to regulate psychiatric-like behaviors.

The GSK3 substrate CRMP2 regulates microtubule dynamics in at least two ways: by binding to tubulin heterodimers to enhance growth at microtubule plus-ends, and by serving as an adaptor between motor proteins and microtubules to promote microtubule elongation.^33,54^ In both cases, phosphorylation of CRMP2 T514 by GSK3 reduces the binding affinity of CRMP2 and destabilizes microtubules. In line with the observed elevation of GSK3 activity in *Ank3*-repressed neuro-2a cells, CRMP2-pT514 level was increased, supporting our hypothesis that changes in microtubule dynamics are mediated through a GSK3ICRMP2 pathway. We were able to substantiate this using a low dose of the CRMP2 antagonist lacosamide to inhibit tubulin polymerization,^56–59^ which prevented lithium and CHIR99021 from rescuing the microtubule changes in *Ank3*-repressed cells. Interestingly, CRMP2 activity has previously been implicated in the lithium responsiveness of BD patients.^22^ In that study, cells derived from lithium-responsive patients had an elevated ratio of CRMP2-pT514 to total CRMP2 (i.e., decreased activity), similar to the elevated ratio we found in *Ank3* repressed neuro-2a cells. This also falls in line with our observation that CRMP2 activity is required for lithium to rescue enhanced microtubule dynamics associated with *Ank3* repression.

Our findings that brain-specific *Ank3* functions in microtubule dynamics advances our understanding of the role of *ANK3* in supporting neuronal function, and potentially its contribution to psychiatric illness. Indeed, accumulating evidence suggests that abnormalities in the cytoskeleton are a potential mechanism for psychiatric illness via impaired microtubule-mediated axonal transport and synaptic plasticity.^60,71–74^ It will be important to investigate how *ANK3* influences microtubule-dependent processes in neurons and whether these processes underlie the association of *ANK3* with psychiatric illness.

## Acknowledgements

The authors wish to thank Dr. Vann Bennett for providing the Ank3 mouse model, Dr. Feng Zhang for providing the sgRNA plasmid, and Francisca Meyer and Vivian Ejsink for advising on RNA sequencing data analysis. We thank Richard Bouley in the MGH Program in Membrane Biology (PMB) Microscopy Core for help with the confocal imaging. The Nikon A1R confocal in the PMB Microscopy Core was purchased using an NIH Shared Instrumentation Grant S10 RR031563-0. Additional support for the PMB Core came from the Boston Area Diabetes and Endocrinology Research Center (DK057521) and the MGH Center for the Study of Inflammatory Bowel Disease (DK043351). This research was supported by a Brain & Behavior Research Foundation Independent Investigator award (TLP) and Young Investigator award (JCG), NIH grants R21 MH099760 (TLP), K22 NS094591 (JCG), and R01 MH107182 (PP), the Massachusetts General Hospital Executive Committee on Research (TLP and JCG), and the European Community’s Seventh Framework Programme (FP7/2007-2013) under grant agreement n° 278948 (TLP). This report reflects only the authors’ views. The funding sponsors are not liable for any use that may be made of the information contained therein.

